# Unconsciously Implanted Visuoauditory Memory in the Presence of Cholecystokinin Retrieved in Behavioral Contexts

**DOI:** 10.1101/241141

**Authors:** Zicong Zhang, Christine Xuejiao Zheng, Yujie Peng, Yiping Guo, Danyi Lu, Xiao Li, Wenjian Sun, Peng Tang, Sherry Ling He, Min Li, Qing Liu, Fuqiang Xu, Gabriel Ng, Xi Chen, Jufang He

## Abstract

We investigated whether visuoauditory association can be artificially implanted in rodents and then retrieved in a behaviorally relevant context. Rats were trained to approach the left or right hole of a behavioral apparatus to retrieve a reward depending on the side of electrical stimulation of the auditory cortex (EAC) they received and mice were fear-conditioned to EAC. Next, an irrelevant visual stimulus (VS) was repeatedly paired with EAC in the presence of cholecystokinin (CCK) or with activation of terminals of entorhinal CCK neurons in the auditory cortex. In subsequent behavioral testing with VS, rats approached the hole associated with reward availability and mice showed a freezing response to the VS. A CCK antagonist blocked the establishment of visuoauditory association, whereas a CCK agonist rescued the deficit of association. Our findings provide a scientific foundation for “memory implantation” and indicate that CCK is the switching chemical for formation of visuoauditory association.

---

The hippocampal system consists of the hippocampus and adjacent entorhinal, perirhinal, and parahippocampal cortices^1^. The entorhinal and perirhinal cortices are the gateway between the hippocampus and the neocortex, and have strong reciprocal connections with the entire neocortex^2,3^. Observations that patients with hippocampal system damage show difficulties forming new long-term memories for facts and events^4,5^ led to our understanding that the hippocampal system is essential for establishing long-term memories^1^. However, patients with hippocampal damage can still recall remote memories^6–8^, suggesting that these memories can be supported by the neocortex^9,10^. Previously, we established a long-term visuoauditory associative memory in rats critically dependent on the entorhinal cortex by pairing a visual stimulus (VS) with electrical stimulation of the auditory cortex^11^.

Many neurons in the entorhinal cortex contain cholecystokinin (CCK)^12–14^. They project to and enable long-term potentiation (LTP) in the auditory cortex by responding to a formerly ineffective sound or light stimulus after pairing with a noise-burst stimulus^15^. Infusion of a CCK antagonist into the auditory cortex prevents the formation of this visuoauditory association, similar to inactivation of the entorhinal cortex^11, 15^.

In this study, we aimed to confirm the formation of visuoauditory associative memory between sound and light stimuli. To test our hypothesis that CCK is a memory-writing chemical for visuoauditory memory in the neocortex, we determined whether the presence of CCK allowed implantation of a memory in the cortex of anesthetized rats retrievable in a behaviorally relevant context. Rats with electrodes implanted in the bilateral auditory cortex were trained to retrieve a reward depending on the auditory cortex stimulated. One hemisphere of the electrically stimulated auditory cortex (EAC) of each rat was infused with CCK and a visual stimulus (VS) paired with stimulation in that hemisphere to establish an artificial link between the VS and EAC. We then tested whether the VS would activate the visuoauditory association so rats would approach the hole that was “engineered” with a reward. We also used CCK-ires-Cre and CCK-CreER (CCK^-/-^) mice to verify our hypothesis. We designed anatomical, physiological, and behavioral experiments to determine characteristics of the CCK neurons. Finally, we performed behavioral experiments to confirm that CCK is the key chemical in cross-modal associative learning.

## Behavioral confirmation of the cross-modal association

To examine whether cross-modal associative memory could be formed and expressed behaviorally using operant conditioning, experimental rats were exposed to combined auditory/visual stimulus (AS-VS) with footshock. Control rats were exposed to VS alone with footshock (Fig.1a). Both groups were trained using the subject-initiated sound-reward protocol (Fig.1a). Rats were required to keep their nose in the central hole for varying intervals of 200-800, 500-1500, and 100-1200ms (stages 3, 4, and 5, respectively). All subjects completed stages 1 and 2 on the first day. The slowest rats required 118, 141, and 149 trials in stages 3, 4, and 5, respectively. The total number of trials in stages 3-5 ranged from 176 to 338. The average reaction time shortened slightly over the course of training (Fig.1b).

**Figure 1.**
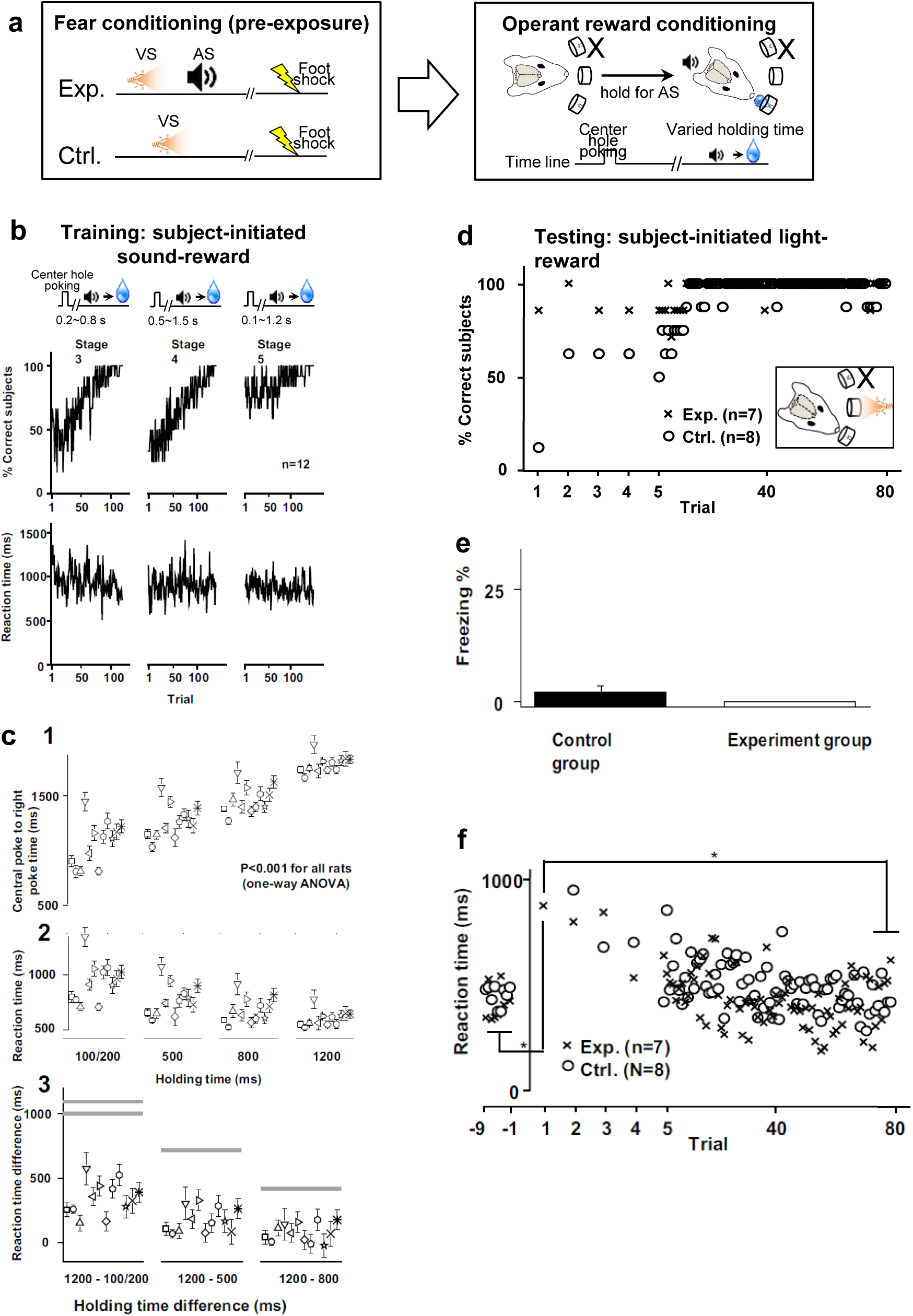
Combining fear and operant reward conditioning to exhibit associative learning between a VS and an AS. **a**. Block diagram showing behavioral protocol combining fear and operant reward conditioning for experimental (Exp) and control (Ctrl) groups. **b**. Top graph shows reward retrieval in stages 3-5 of the subject-initiated sound-reward protocol. The middle and bottom panel shows the success rate and the reaction time (the time to retrieve the reward after the sound was triggered in stages 3-5 for all rats. **c**. Detailed data show the subjects’ responses in stage 5. **1**. The interval between nose-poking in the central hole and moving to the rightmost hole, shown separately for each holding time (100-ms and 200-ms holding times were pooled). Each data point represents an individual rat. The interval differed significantly across holding times (P<0.001, one-way ANOVA). **2**. Time interval before reaching the rightmost hole for the reward after the sound was triggered, shown separately for each holding time. **3**. Differences in the reaction times associated with different holding times. Bars indicate the time difference between the two holding times in question. **d**. Percent of successful trials in the experimental group (crosses) and control group (open circles) in test trials in which the AS was replaced by the VS (i.e., SiLR protocol). **e**. Percent time spent freezing during the 60-s period after the VS was presented. **f**. Time to retrieve the reward after the sound (trial<0) and the light (trial>0) was triggered in the experimental and control groups. Only rats that successfully retrieved the reward were included in the analyses. Values from trial 1 vs. the mean values on trials 72-80 and vs. the mean values on the last nine subject-initiated sound-reward trials were compared (F=11.64, P<0.001, one-way ANOVA; Post hoc, *P<0.001).

There was overlap between the earliest and latest reward windows. If a cue was delivered at the shortest (100ms) or longest (1200ms), a rat could retrieve the reward at any point between 100-2100ms or 1200-3200ms, respectively. A rat could solve the task by simply withholding responding for ~1500ms, then entering the right hole and retrieving the reward, and the decrease in reaction times associated with lengthened holding time suggests that the rats may have (Fig.1c2). On the other hand, there were significant differences in all rats in the average interval between nose-poking in the central hole and moving to the rightmost hole (Fig.1c1). The difference between the reaction times on trials involving different holding times was far below the difference in the holding times (Fig.1c3), suggesting that the decrease in the reaction time as the holding time lengthened was a function of the longer time to prepare for reward retrieval while the rat was waiting for the cue.

The reaction time to retrieve the reward after the AS was triggered indicated that the rats used the AS as a signal of reward availability (Video1, 3; Fig.1c3). We thus replaced the AS with the VS. 6 of 7 rats in the experimental group retrieved the reward on the first trial, and all rats did on the second trial. In contrast, 7 of 8 rats in the control group failed to retrieve the reward on the first trial (Fig.1d). These results suggest that the experimental group successfully associated the AS and VS and used the VS to recall the AS before retrieving the reward (Fig.1d). The purpose of the control group was to determine whether VS-triggered reward-reception behavior in the experimental group was based on procedural sequence knowledge (i.e., first nose-poke into the central hole for initiation, then wait for any sort of cue, then retrieve the reward). The differences in these groups indicates that the experimental group did use visuoauditory associative memory to complete the task.

In both groups, conditioned fear in the presence of the VS was extinguished after the rats completed the subject-initiated sound-reward protocol and the subject-initiated VS-reward protocol (Fig.1e). This result indicates that the deficit in reward reception seen in the control group was not due to fear of the VS.

The rats of the experimental group initially paused, then reacted quickly to retrieve the reward after replacement of the AS with the VS (Video2, 4; Fig.1f). Their reaction time was significantly longer on the first trial with the VS than on the earlier trials with the AS (Fig.1f), which may reflect the time needed for recalling the auditory cue from the visual cue. The reaction time with the VS, however, decreased over time. A similar trend was observed in the control group, indicating learning of new association between the VS and reward.

## Auditory cortex stimulation as cue for reward

To introduce more options for the animals to potentially create designated associative memory, we utilized 3 holes in the water reward system and adopted a two-alternative forced choice (2AFC) auditory task. Rats were trained to retrieve the water reward from the left or right hole of the apparatus depending on whether the right or left auditory cortex was stimulated by implanted electrodes.

Electrode arrays and guide cannulae were implanted in the bilateral auditory cortex of each rat (Fig.2a). The rat was initially trained to nose-poke in the center hole of a behavioral apparatus to trigger AS of high or low frequency indicating water reward from left or right hole (Fig.2b; Video 5). In the next stage, AS was replaced with electrical stimulation of the left or right auditory cortex (EACL or EACR, respectively; Fig.2b). EACR indicated the availability of a water reward in the left hole, and vice versa (Video 6). Across 4 stages of training (Fig.2b), 10 rats reached a success rate of 90% at the final stage (Fig.2c), indicating they successfully learned to utilize EAC as a cue to approach the correct hole for reward retrieval.

**Figure 2.**
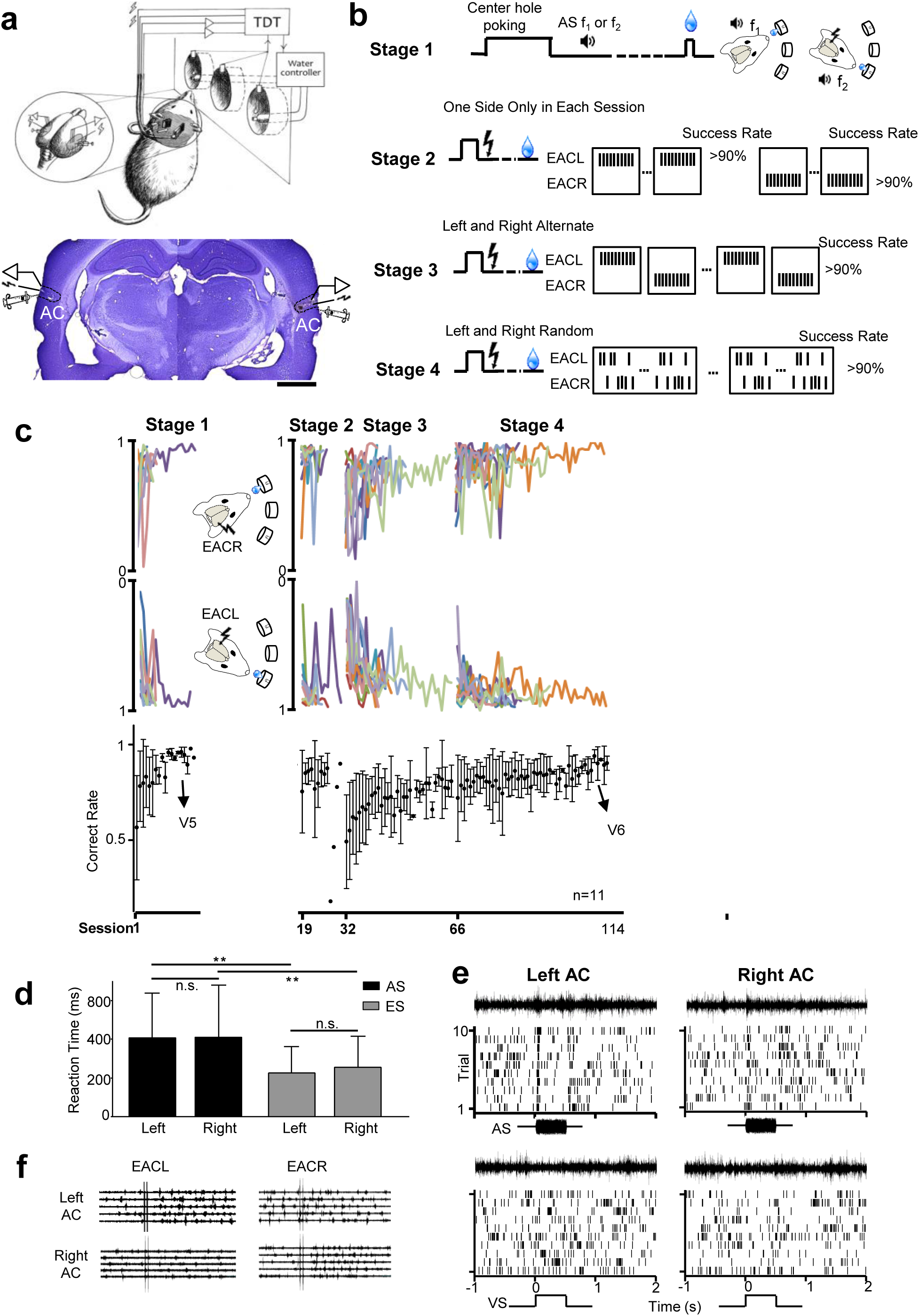
EAC as a cue for reward availability. **a**. Upper panel: Left, center, and right holes of a behavioral apparatus. Lower panel: Nissl staining shows the locations of the bilateral implantations of electrodes and drug infusion cannulae. **b**. Diagram of training protocol for stages 1-4 and timing of nose-poke detection in the center hole, cue presentation, and reward delivery. During stage 1, rats learned to use two different sounds as cues for reward availability in the right or left hole. During stage 2, rats learned to use EACL or EACR as a cue for reward availability in the right or left hole, respectively. Hemispheres were alternately stimulated between sessions, with 100 trials per session. During stage 3, hemispheres were alternately stimulated between sessions, with 10 trials per session. During stage 4, hemispheres were stimulated in a pseudo-random manner in individual trials. **c**. Success rate in reward retrieval across four training stages for the left hole (upper panel) and the right hole (middle panel). The correct rat for the training progress over both sides across four training stages is shown in the lower panel. Video 5 (V5) and V6 demonstrate the training process. **d**. Means and standard deviations (SDs) for behavioral reaction time, defined as latency from the onset of the AS or EAC to leaving of the center hole. \*\**p*<0.001. **e**. Raster plots show neuronal responses in the left and right auditory cortex (AC) to the AS (upper panels) and the VS (lower panels). **f**. Raw traces show neuronal responses in the left and right AC to EACL and EACR.

There were no significant differences between hemispheres in behavioral reaction times to the cue, regardless of whether the cue was sound or EAC (Fig.2d). Rats showed shorter reaction times when the cue was EAC than AS (Fig.2d). This reduction in reaction time is likely because of better learning during advanced training stages and/or shortened stimulation perception, with direct EAC being faster than an AS traveling from the cochlea to the auditory cortex.

Neuronal responses to light and sound stimuli were recorded from both hemispheres before implantation of the electrodes. Neurons in both hemispheres responded to a noise-burst stimulus, but showed no or weak responses to the VS (Fig.2e). EACL affected neuronal responses in the ipsilateral hemisphere (Fig.2f left, excitatory response in upper panel), but not in the contralateral hemisphere (Fig.2f left, lower panel), and vice versa (Fig.2f right).

## Retrieval of unconsciously implanted memory in a behaviorally relevant context

We then examined whether the presence of CCK would allow us to implant a visuoauditory associative memory in the auditory cortex of anesthetized rats that could be retrieved in a behaviorally relevant context. After learning to use EAC as a cue for reward availability, rats underwent baseline testing during which EAC was sometimes replaced with the VS (a light flash) (Fig.3a). Typically, 10 VSs were randomly inserted into a total of 1000 training trials. Rats usually showed no fixed response to the VS in terms of approaching a certain direction (Fig.3b, Supplementary Fig.1). Rats 1 and 2 showed a preferred direction, though rats 3-5 showed no preference (Videos 7 and 9; Fig.3b). After baseline testing, VS was paired with EAC of the non-preferred hemisphere and CCK infusion (Fig.3b). Thus, the “engineered” direction was opposite to the preferred direction. Rats then underwent post-intervention testing following identical procedures as baseline testing for up to 4 weeks. Importantly, no reward was delivered after the VS to prevent an independent association between light and reward.

**Figure 3.**
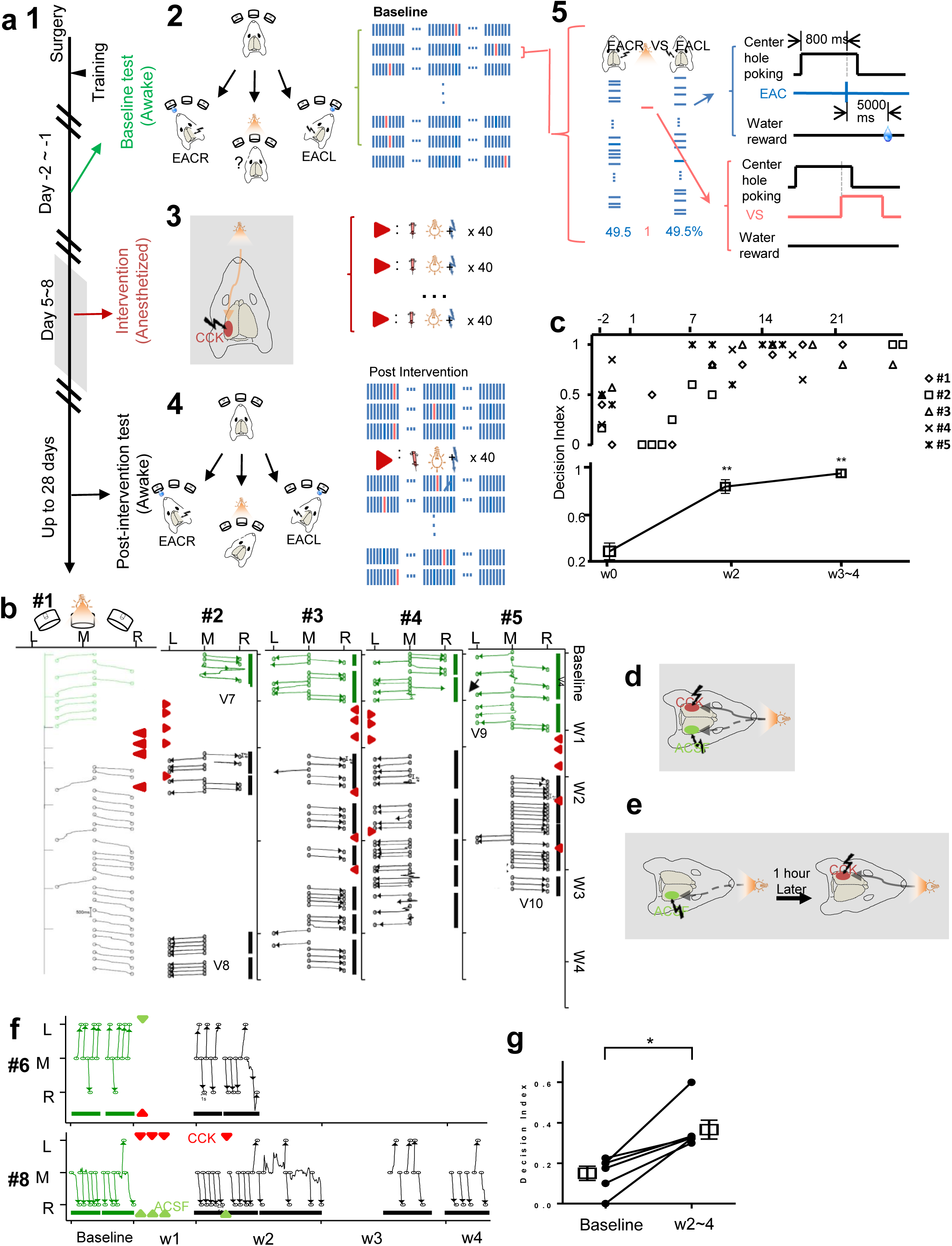
Behavioral read-out of memory implanted under anesthesia. **a**. Diagram of training protocol. After rats were trained to use cortical stimulation as a cue for reward availability, they underwent 3 experimental phases: baseline testing while awake, memory implantation under anesthesia and in the presence of CCK, and post-intervention testing while awake. **1**. The timeline for the entire experiment. **2**. During baseline testing, a VS occasionally replaced EAC, and behavioral responses to the light (marked by “?”) were recorded. **3**. During the memory implantation phase, rats were under anesthesia and CCK was infused into the auditory cortex of one hemisphere before pairings of the VS (light flash) with EAC of that hemisphere (EACL, in the figure). Stimulus pairings occurred for 40 trials per day on multiple days. **4**. Post-intervention testing was identical to baseline testing. **5**. Presentation of the EAC (blue) and VS (red) during baseline and post-intervention test. **b**. Behavioral responses (trajectory of head movement) to the VS during baseline (green curves) and post-intervention (black curves) testing of rat #1-5. ‘L’ indicates left hole approach, ‘R’ indicates right hole approach, and ‘M’ indicates the middle hole. Stimulus pairings after CCK infusion are marked by red arrowheads. V7 and V9 demonstrate behavioral responses before stimulus pairings. V8 and V10 demonstrate behavioral responses after stimulus pairings. **c**. Individual values (upper) and means (lower) for Decision Index across days. Decision Index was defined as ‘1’ when the rat approached the “engineered” hole, ‘0’ when it approached the other hole, and ‘0.5’ when it approached neither hole. \*\**p*<0.0001, one-way ANOVA with post-hoc Dunnett’s multiple comparison tests. **d**. Schematic drawing shows the preparation of the infusion of CCK and artificial cerebrospinal fluid (ACSF) and pairing of the VS and EAC. CCK and ACSF were simultaneously infused into the left and right hemispheres, respectively, prior to pairings of the VS and EAC of both hemispheres. **e**. Schematic drawing shows the preparation of sequential infusion of ACSF and CCK. ACSF and CCK were infused into different hemispheres within the same period of anesthesia. Stimulus pairings occurred after each infusion. **f**. Behavioral responses to the VS presentations of 2 exemplary rats. Arrowheads indicate stimulus pairing sessions followed by infusion of CCK (red) or ACSF (green). **g**. Decision Index before (w0) and after stimulus pairings (w2-4). n=6, *p<0.05.

During baseline testing, rat #1 tended to approach the left hole in response to the VS (Fig.3b). In week 1, the VS was paired with EACL under anesthesia and CCK infusion to “engineer” approaches to the right hole (Fig.3b). On day 9, the rat approached the right “engineered” hole in 4 of 5 trials. On days 22 and 28, the rat approached only the right hole in response to VS.

For rat #2, the VS was paired with EACR and CCK infusion for 5 sessions in weeks 1 and 2. During post-intervention testing, the rat approached the right hole in response to the VS on the first few days, then started to approach the left “engineered” hole later. On days 27 and 28, the rat approached the left hole in all trials (Video 8). For rats #3-5, stimulus pairings occurred between days 1 and 5, and post-intervention testing started on day 7 and onward (Video 10 for rat #5). All rats consistently approached the “engineered” hole in response to the VS (Fig.3b).

These “engineered” changes in behavior were maintained for several days without further intervention. Behavioral changes were maintained at least 10, 20, 12, 7, and 4 days, respectively, for rats #1, #2, #3, #4, and #5. A Decision Index scoring system was created to quantify each animal’s directional decision upon VS. A score of 1 was assigned when animal approached the engineered hole, and 0 for the opposite hole. The Decision Index increased significantly over time (Fig.3c). Together, these results demonstrate that the novel VS was associated with EAC of the hemisphere with CCK, and this association subsequently guided the animal to the corresponding hole for water reward.

## Controlling for non-specific effects of infusion and stimulus pairings

We conducted two control experiments to rule out the possibility that the changes of animals’ behavior were due to non-specific factors (e.g., anesthesia, drug infusion, researcher’s handling): (1) simultaneous infusion of CCK into one hemisphere and artificial cerebrospinal fluid (ACSF) into the other hemisphere with simultaneous pairing of the VS with EAC of both hemispheres (Fig.3d), (2) infusion of ACSF and stimulus pairings in one hemisphere followed 1h later by infusion of CCK and stimulus pairings in the other hemisphere (Fig.3e). All 6 rats in both experiments showed an increase in Decision Index during the 2 weeks after stimulus pairings (see Fig.3f; Decision Index from 0.15 ± 0.03 to 0.37 ± 0.05, *p*<0.05, paired t-test, Fig.3g). These results suggest that the association was established specifically via the function of CCK.

## Responses of auditory cortex neurons to the VS after unconscious memory implantation

Underlying the associative memory reflected in the behavioral study, neural plasticity should occur at the cellular level. Therefore, we monitored the neural activity of the auditory neurons in response to VS during CCK infusion and stimulus pairing. An example neuron in an anesthetized rat did not respond to the VS before stimulus pairings (Day 1) but gradually started to respond to VS after stimulus pairings (Days 2-17, Fig.4a). The mean Z-score of neuronal responses significantly increased after stimulus pairings (Fig.4b). Next, we examined whether the altered neuronal plasticity observed in anesthetized rats could also be observed in awake rats. An example neuron did not respond to the VS before conditioning but showed a response to the VS after intervention (Fig.4c). The mean Z-score increased significantly after stimulus pairings (Fig.4d).

**Figure 4.**
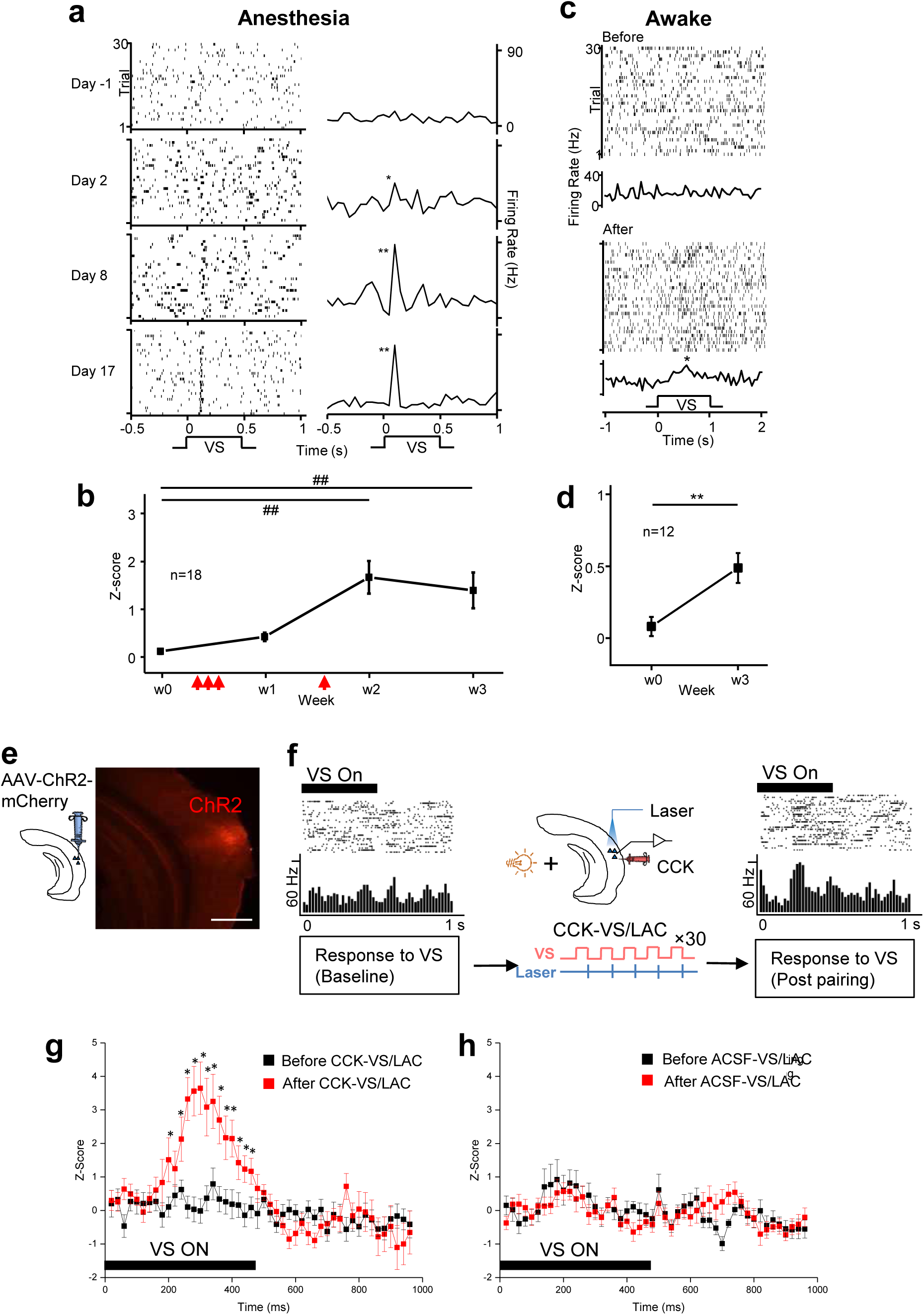
Changes in response of auditory cortical neurons to the VS. **a**. Raster plots (left) and peri-stimulus time histograms (PSTHs, right) showing neuronal responses to the VS before (Day – 1) and after (Days 2, 8, 17) CCK infusion and stimulus pairings of VS/EAC. Data were collected from the anesthetized rat. **b**. Z-scores were calculated based on differences between average neuronal firing during a 150-ms period after the VS and an equivalent period of spontaneous firing. Data were collected from anesthetized rats. **c**. Neuronal responses to the VS before and after stimulus pairings. Data were collected from the behaving rat. **d**. Z-scores were calculated based on differences between average neuronal firing during a 200-ms period after the VS and an equivalent period of spontaneous firing. ##*p*<0.01, ANOVA; \**p*<0.05, \*\**p*<0.01, paired *t*-test. **e**. Schematic drawings for positions of virus injection in the auditory cortex (left) and virus expression in the auditory cortex. **f**. Raster plots and PSTHs show neuronal responses to the VS (light flash) before (leftmost) and after (rightmost) CCK infusion and stimulus pairings of VS/Laser stimulation of the auditory cortex (CCK-VS/LAC). Middle panel shows the schematic drawing of the preparation. Lower panel shows the procedure of the pairing. Data were collected from the anesthetized rat. **g-h**. Z-scores show that the neuronal responses to the VS before and after the CCK-VS/LAC (**g**) or the ACSF-VS/LAC (**h**). Z-scores were calculated based on differences between average neuronal firing on 20-ms time bins after the VS and an equivalent period of spontaneous firing. n=8, *p<0.05, two-way ANOVA.

We then questioned what type of neurons are involved in such neural plasticity and account for visuoauditory association. Given that excitatory cortical neurons are usually involved in neuroplasticity, we induced expression of the light-evoked ion channel channorhadopsin-2 (ChR2) into the excitatory auditory neurons, and implanted optic fiber and recording electrodes in the auditory cortex to manipulate and record these neurons (Fig.4e, Supplementary Fig.2). We examined whether pairing the VS with activation of these neurons in the presence of CCK would change their response to the VS. The auditory neuron response to the VS changed after 30 trials preceded by CCK infusion (Fig.4f; Fig.4g, 3.56±0.74 vs 0.03±0.37, Z-score, p<0.01), whereas nothing changed after the same pairing preceded with ACSF infusion (Fig.4h; -0.04±0.23 vs 0.11±0.32, Z-score; p>0.05). In the presence of CCK, pairing the VS and activation of the auditory neurons facilitated neuronal responses to the VS in the auditory cortex.

## High-frequency (HF) stimulation of terminals of entorhinal CCK neurons in the auditory cortex enabled the potentiation of neuronal response to the VS

To verify our hypotheses that high-frequency (HF) stimulation is sufficient to trigger entorhinal CCK neurons to release CCK on their terminals in the auditory cortex and the released enable auditory neurons to respond to VS after VS/EAC pairing, we induced ChR2 expression in entorhinal neurons driven by Cre recombinase in CCK-ires-Cre and CCK-CreER (CCK^-/-^) mice. We implanted optic fiber and electrode arrays in the auditory cortex for both stimulation and recording (Fig.5a). CCK neurons expressing ChR2 were labeled in the entorhinal cortex of both mice (Figs. 5b-c, g-h). Their terminals were found in both auditory (Figs. 5d-e, i-j) and visual cortices (Figs. 5d, f, i, k). Field excitatory postsynaptic potential (fEPSP) evoked by laser stimulation of a single pulse or an 80-Hz pulse train toward terminals in the auditory cortex was found in both CCK-ires-Cre (Figs. 5l-m) and CCK^-/-^ (Figs. 5r-s) mice, indicating that entorhinal CCK neurons that project to the auditory cortex were mainly excitatory. Neurons in the auditory cortex of both mouse types showed responses to the AS, as shown in fEPSP (Figs. 5n, t) and unit responses (Fig.5p, v). Moreover, we observed small fEPSP and unit responses to the VS in the auditory cortex of CCK-ires-Cre mouse (Figs. 5o, q), but no response in the auditory cortex of CCK^-/-^ mouse (Figs. 5u, w). The lack of visual response in the auditory cortex of the CCK^-/-^ mouse might reflect a deficit in cross-modal association without CCK.

**Figure 5.**
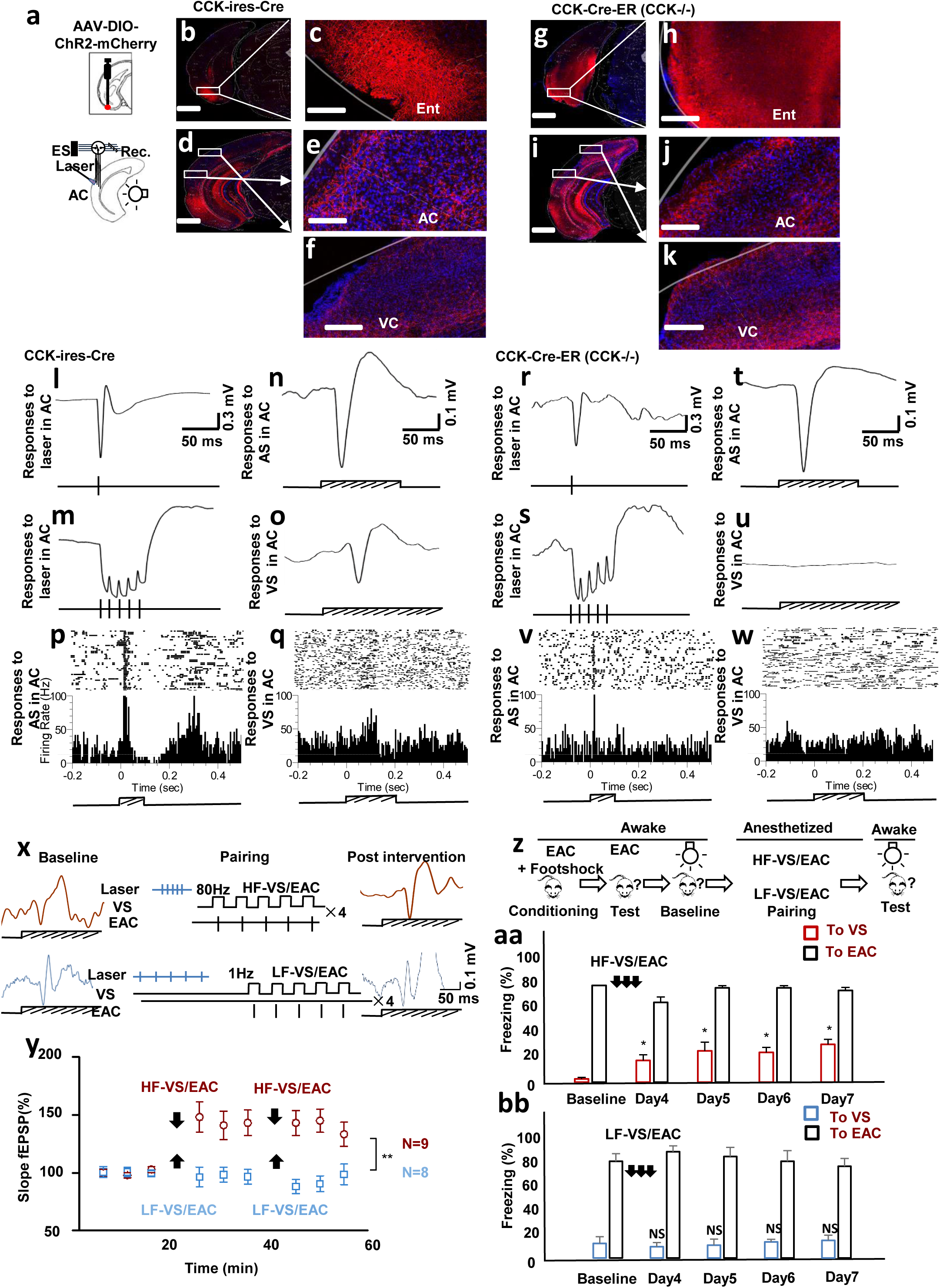
High-frequency activation of CCK-containing entorhino-neocortical projections enables the association between the VS and EAC, leading to behavioral changes. **a**. Schematic drawings for positions of virus injection in the entorhinal cortex and implantations of the laser fiber and stimulating/recording electrodes in the auditory cortex (AC). **b**. AAV-DIO-ChR2-mCherry was injected into the entorhinal cortex of CCK-ires-Cre mice. Images show virus expression in the entorhinal cortex (**b-c**), the AC (**d-e**), and the visual cortex (VC) (**d**, **f**). **g**. AAV-DIO-ChR2-mCherry was injected into the entorhinal cortex CCK-CreER (CCK^-/-^) mice. Images show virus expression in the entorhinal cortex (**g-h**), the AC (**i-j**), and the VC (**i**, **k**). Scale bars: 500µm for **b**, **d**, **g**, **i**; 100 µm for **c**, **e**, **f**, **h**, **j**, and **k**. **l-o**. fEPSPs to laser stimulation in the AC in single pulse (**l**) and in pulse train (**m**), AS(**n**), and VS (**o**) in the AC of CCK-ires-Cre mice. **p-q**. Raster plots and PSTHs show unit responses to the AS and VS in the AC of CCK-ires-Cre mice. **r-u**. fEPSPs to laser stimulation in the AC in single pulse (**r**) and in pulse train (**s**), AS (**t**), and VS (**u**) in the AC of CCK^-/-^ mice. **v-w**. Raster plot and PSTH show unit responses to the AS and VS in the AC of CCK^-/-^ mice. **x**. Representative fEPSPs before (left) and after (right) the HF-VS/EAC and LF-VS/EAC protocols (middle). Upper panel shows fEPSPs and the protocol of HF-VS/EAC, and the lower panel shows fEPSPs and the protocol of LF-VS/EAC. **y**. Normalized slopes of fEPSPs after the HF-VS/EAC (red circle) or LF-VS/EAC (blue square) pairing protocol. **p<0.001, two-way ANOVA. **z**. Cued fear conditioning and stimulation protocols. **aa-bb**. Bar charts show the percentage (mean±SEM) of time spent freezing in response to the conditioned EAC and the paired VS before and after the HF-VS/EAC (**aa**) or LF-VS/EAC (**bb**) pairing. *p<0.05, NS, Not significant, one-way ANOVA.

Next, we adopted a pairing paradigm (Fig.5x) in which HF laser activation of entorhinal CCK neuron terminals was followed by 5 paired presentations of the VS and EAC. The pairing paradigm (referred to as HF-VS/EAC) was repeated 4 times. CCK^-/-^ mice were not included in the control group because neurons in their auditory cortex showed no response to the VS. We utilized the same mouse type and similar paradigm but with low-frequency (LF) laser activation (LF-VS/EAC) of terminals of entorhinal CCK neurons in the auditory cortex for the control (Fig.5x). The neuronal response of fEPSP (amplitude and slope) to the VS increased after the HF-VS/EAC (Fig.5x; Fig.5y, 140.1±6.91%, F=18.746, p<0.001). There were no changes in amplitude and slope of the fEPSP after the LF-VS/EAC pairing (Fig.5x; Fig.5y, 99.5±3.97%, F=0.322, p=0.725). There was a significant difference in change of fEPSP between the experimental and control groups at the end of the recording after two pairing (Fig.5y, 139.6, vs 96.4%, F=44.949, p<0.001), indicating that the potentiation in the neuronal response to the VS induced with HF-VS/EAC, but not with LF-VS/EAC.

## Changes in the neuronal response during anesthesia reflected in behavioral test

We hypothesized that the visuoauditory association of enhanced neuronal response to the VS during the above paradigm in anesthetized mice should be reflected in behavioral experiments, so we pre-conditioned the EAC with footshock for 3 days before pairings and compared the behavioral response to the VS after the pairing with that before the pairing (Fig.5z). Mice showed significantly increased freezing responses to the VS after HF-VS/EAC (Video11-14; Fig.5aa; 3.33±1.78% before vs 22.5±6.29% at day 4, p<0.05, F=4.986), but not after LF-VS/EAC (Video15-18; Fig.5bb; 11.7±2.44% before vs 10.0±2.18% at day 4, p=0.791, F=0.423). The fear response to the VS was maintained at day 7, or 4 days after the last HF-VS/EAC pairing (Fig.5aa; 3.33±1.78% before vs 39.2±5.55% at day 7, F=4.986, p<0.05).

## Application of CCK antagonist blocked the establishment of visuoauditory association

To test whether CCK (which we had previously tested artificially) is necessary for the establishment of natural associative memory, we examined the formation of associative memory with a fear response test. First, CCKB receptor antagonist or ACSF was injected bilaterally into the auditory cortex, followed by pairing of VS with AS. On day 4, baseline tests for freezing response to the AS and VS were performed for 3 trials before fear conditioning. After fear conditioning, freezing responses to the AS and VS were further examined (Fig.6a). Understandably, mice showed no freezing response to the AS before the conditioning, but a high freezing rate to the AS after the conditioning (Video21, 22, 25, 26; Fig.6b; 7.594±3.77% vs 68.7±4.30%, F=150.816, p<0.001). The ACSF mice group also showed significantly increased freezing response to the VS, indicating an association between the AS and VS was established during the pairings (Video23, 24; Fig.6b; 4.07±0.92% vs 24.4±3.77%, pre- and post-intervention of ACSF group; p<0.001, F=27.563). However, the infusion of L-365,260 into the bilateral auditory cortex blocked this association, resulting in a nil response to the VS (Video19, 20; Fig.6b; 4.27±1.23% vs 2.50±1.64%, pre- and post-intervention of L-365,260 group; F=0.746, p=0.402). Mice showed a promising association between the AS and footshock, as indicated by high freezing rate to the AS. An association between the VS and AS was not established. There was a significant difference between the freezing rates of experimental and control groups (Fig.6b; 2.50±1.64% vs 24.4±3.77%, respectively; F=26.098, p<0.001). These results demonstrate that the CCKB receptor antagonist prevented the establishment of an association between the VS and AS, and suggest an essential function of CCK in visuoauditory association formation.

**Figure 6.**
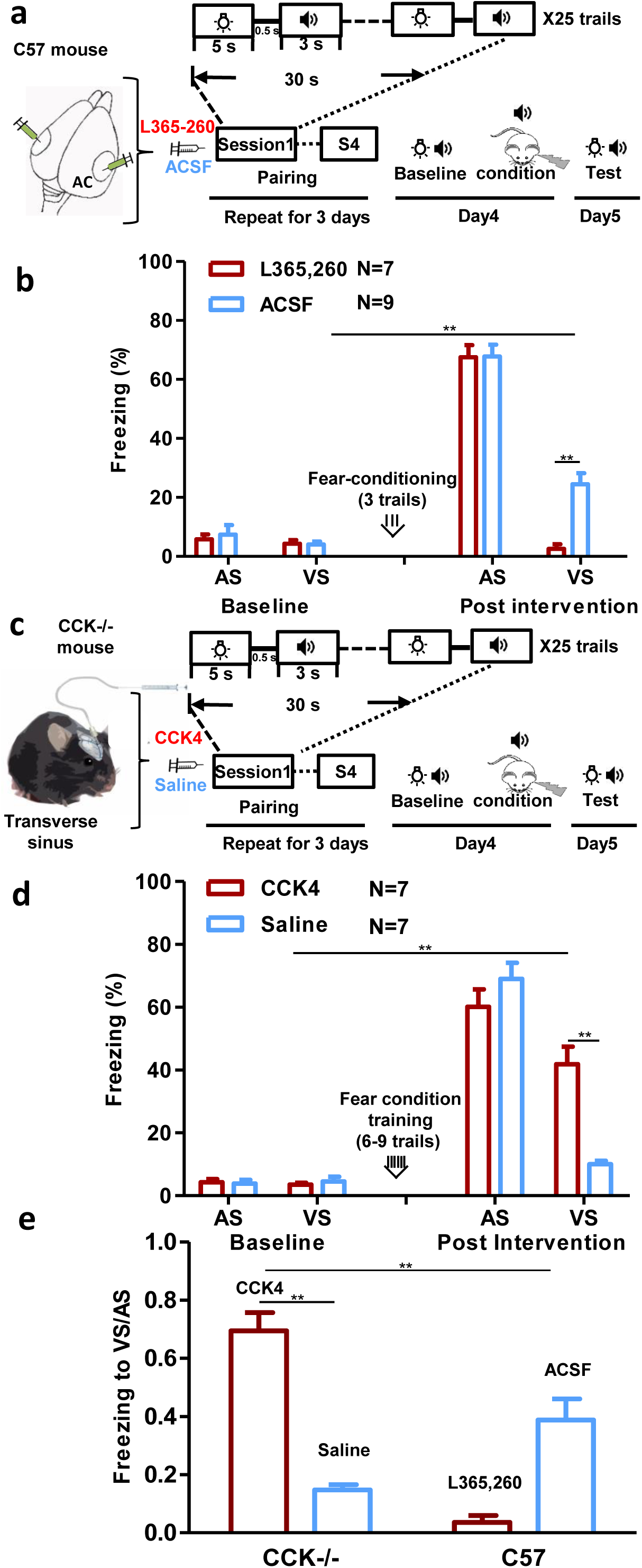
Visuoauditory associative memory could not be formed without CCK. **a-b** Formation of visuoauditory associative memory blocked by CCKB receptor antagonist in the AC. **a**. Diagram of training protocol for wild-type mice to associate the VS and AS. **b**. Bar chart showing freezing percentages to the AS and VS before and after the conditioning. **p<0.01, one-way ANOVA, post-hoc Tukey test. **c-d**. Formation of visuoauditory associative memory was rescued in CCK^-/-^ mice after intravenous administration of CCK4. **c**. Diagram of implanted drug infusion cannula. The training protocol for mice to associate the VS and AS was similar to **a**, but with systemic administration of CCK-4 through the implanted drug infusion cannula in the transverse sinus. **d**. Bar chart showing freezing percentages of CCK^-/-^ mice to the VS and AS before and after the intervention (drug infusion before the VS/AS pairing). **e**. The ratio (mean±SEM) of freezing in response to the VS over that to the AS after the intervention of wild-type and CCK^-/-^ in all conditions are summarized in the bar chart. **p<0.01, one-way ANOVA.

## Systemic administration of tetropeptide CCK rescued the formation of associative memory in CCK^-/-^ mice

Like the CCK antagonist group, we expected that CCK^-/-^ mice would show a deficit in the formation of associative memory, which we tested (Fig.6c). With some preliminary study, we understood that the AS could be conditioned with a delay fear conditioning in CCK^-/-^ mice. Notably, 6-9 trials were needed for CCK^-/-^ mice to produce the freezing rate of >60% in response to the conditioned AS, whereas only 3 trials were needed for wild-type mice (data not shown), suggesting a general associative learning deficit in the CCK^-/-^ mice. The CCK^-/-^ mice in the control group (with saline injection) consistently showed minimal freezing response to VS after visuoauditory association and fear conditioning (Video31-34; Fig.6d). To determine whether systematic administration of CCK could rescue this deficit, we administrated a CCK agonist (CCK-4) that can penetrate the blood-brain barrier through an implanted drug infusion cannula into the transverse sinus. The experimental group with CCK-4 injection showed a significantly higher freezing rate compared to the control (Video27-30; Fig.6d; 43.8±6.16% vs 10.0±1.13%; F=34.269, p<0.001), indicating that the visuoauditory association was rescued upon CCK-4 administration.

To better compare the strength of visuoauditory association in different experimental conditions, we calculated the ratio of the freezing response to the VS compared with that to the AS after conditioning (Fig.6e). The ratio of the L-365,260 group was lowest among all groups, demonstrating a nearly complete abolishment of the visuoauditory association. Interestingly, the ratio of the CCK-4 group was the highest among all groups (62.2%±9.07% vs 15.3%±2.04% saline in CCK^-/-^ mice, p<0.001, F=16.114; vs 38.8%±7.26% ACSF in wild-type mice, p<0.01). This result indicates the overexpression of CCK receptors in CCK^-/-^ mice, leading to the highest association between the VS and AS.

## DISCUSSION

Infusion of CCK into the auditory cortex of anesthetized rats enabled a neuroplastic change that caused auditory neurons to start responding to a VS after the light was paired with EAC. This artificial visuoauditory association lasted for at least 20 days. Not only did auditory cortex neurons respond to the VS in awake rats, but this artificial association between the VS and activation of the auditory cortex formed under anesthesia was translated into behavioral action, providing a scientific foundation for “memory implantation.”

Pairing the VS and the activation of the infected neurons in the auditory cortex in the presence of CCK changed neuronal response to the VS. HF activation of the terminals of the entorhinal CCK neurons in the auditory cortex, followed by pairing the VS and EAC, enhanced neuronal responses to the VS in the auditory cortex of anesthetized mice. When the mice were pre-conditioned EAC with footshock, they showed significantly increased freezing responses to the VS after the pairings. The fear response to the VS was maintained for 4 days after the HF/VS/ES pairing. Application of CCK antagonist blocked the establishment of visuoauditory association reflected in fear response tests, in which the VS was paired with the AS immediately after the injection of CCK antagonist into the auditory cortex. When fear was only conditioned with AS, mice showed a fear response to VS in the control group, but not the CCK antagonist group. CCK^-/-^ mice had a deficit in forming the visuoauditory association with the above pairing paradigm, but systemic administration of CCK-4 rescued the formation of associative memory in the CCK^-/-^ mice.

In a previous study, we established an artificial association between a VS and EAC through classical conditioning, such that auditory cortex neurons began responding to a VS^11^. Inactivation of the entorhinal cortex blocked the establishment of this association. Previous experiments in rats demonstrate that inactivation of the perirhinal cortex disrupts the encoding, retrieval, and consolidation of object recognition memory^16^, whereas deep-brain stimulation of the medial temporal cortex in humans boosts memory performance^17^. Together, these findings add to our current understanding that the formation of associative memories requires interactions between the neocortex and the hippocampal system^4,5^.

CCK is the most abundant of all neuropeptides^18^, and heavy labeling of CCK-containing neurons is observed in perirhinal and entorhinal cortices^12^. Recently, we found that retrogradely labeled neurons in the entorhinal cortex after infusion of a tracer in the auditory cortex are mostly CCK-containing neurons. Moreover, activation of the entorhinal cortex enabled neuronal plasticity in the auditory cortex, but this was minimized by local infusion of CCK antagonist into the auditory cortex^15^. Therefore, we propose that the hippocampal system sends a memory-encoding signal, the neuromodulator CCK, to the neocortex to enable the formation of a new associative memory through its entorhinal/perirhinal gateway. On the other hand, CCK facilitates neuronal plasticity in hippocampal neurons *in vitro*^19^. Further support for our hypothesis that CCK plays a critical role in behavior, are studies showing that blockade of CCK receptors suppresses conditioned fear^20^, and deletion of the CCKB receptor gene reduces anxiety-like behavior^21^. Furthermore, activation of CCKB receptors in the amygdala potentiates the acoustic startle response^22,23^, and application of a CCKB receptor antagonist attenuates fear-potentiated startle^24^. Findings that mice lacking the CCK gene exhibit poor performance in a passive avoidance task and impaired spatial memory^25^ could be interpreted as the result of a memory-encoding deficit in the brain.

Using fear conditioning and optogenetic techniques in rats, memory-encoding neurons created a false association between a particular context and footshock that was never delivered^26^. Similarly, we implanted an artificial association between a VS and an EAC while rats were under anesthesia and CCK infusion. Subsequently, the rats used an existing association between EAC and a water reward and thus used the VS to guide their retrieval of the reward. Although the visuoauditory associative memory implanted in the auditory cortex was quickly reflected in neuronal responses to the VS, it took several days before the association was reflected in the rats’ behavior. This delay may be due to the time needed for the newly implanted memory in the neocortex to be registered in the hippocampus^27^ or to be linked with place cells in the hippocampus for directional guidance^28^.

Deep-brain stimulation of the entorhinal cortex while subjects learned locations of landmarks enhanced their subsequent memory of these locations^17^. Evidence suggests that coordinated oscillation of entorhinal and hippocampal neurons in 20-40 Hz links to the encoding and retrieval of the associative memory^29^, whereas gamma-band (40-120 Hz) oscillation links to cortical plasticity and formation of new memories^30,31^. However, multimodal associative memory is likely based on the association of multiple cortical areas^32,33^. Formation of paired visual associations or paired visuoauditory associations critically depends on the perirhinal and entorhinal cortex^11,34^. In the present study we confirm that HF activation, but not LF stimulation, of the neocortical projection terminals of entorhinal neurons leads to LTP in fEPSP to the VS in the auditory cortex. Modifying the synaptic strength leads to memory^35^. The potentiated neuronal responses to the VS led to an increased freezing time when the VS was presented, indicating that association between the AS and ES in the auditory cortex was strengthened. Given that we used CCK-ires-Cre mice, neurons infected with AAV-DIO-ChR2-mCherry in the entorhinal cortex were CCK neurons. Earlier studies indicate that release of neuropeptides occurs slowly in response to repetitive firing^36,37^.

Consistent with previous findings that CCK is associated with learning and memory^20,24^, we found that CCK^-/-^ mice show deficits in associating the visual and auditory stimuli. CCK^-/-^ mice needed 1-2 times more conditioning trials to associate the sound cue with footshock (even with delayed conditioning protocol). In the present study, CCK-containing entorhinal neurons in CCK-ires-Cre mice labeled with AAV-DIO-ChR2-mCherry projected all the way to the auditory cortex and visual cortex. Laser stimulation of these terminals elicited fEPSPs in the auditory cortex, suggesting that they are glutamatergic neurons. Association of the artificial manipulation using electrical stimulation and natural stimulus suggests potential neuroengineering and therapeutic applications. The rescuing effect of intravenous application of CCK4 in forming associations between the VS and AS in the brain further proved that the CCK is the key chemical for encoding of visuoauditory associative memory in the brain.

These results, together with our 2 recent studies^11,15^, indicate that the entorhinal cortex is important for the establishment of visuoauditory associative memory, and that CCK is involved in this bridging process. In the presence of CCK, auditory cortex neurons begin to respond to a VS after pairings of the VS and ES of the auditory cortex. Moreover, this altered neuronal responding can be interpreted as an artificial memory in a behaviorally relevant context. Although we have more questions about the temporal gap between stimulus pairings and behavioral changes, our study provides a new perspective on how memory is written in the neocortex. Furthermore, it demonstrates that an artificial memory established in the neocortex can be translated to behavioral action, providing a scientific foundation for “memory implantation.” These results also provide insight for future efforts to rewire malfunctioning brains to bypass lost functions.

## ACKNOWLEDGEMENTS

The authors thank Wendy Zhou for graphic drawings and Eduardo Lau for administrative and technical assistance. This work was supported by Hong Kong Research Grants Council, National Key Basic Research Program of China, Natural Science Foundation of China, and Health and Medical Research Fund (2013CB530900, 561212M, 561313M, 11101215M, 11166316M, 11102417M, C1014-15G, T13-607/12R, 31200852, 03141196, 01121906, 31171060, 31371114, 31571096, 31671102). We also thank the following charitable foundations for their generous support: Charlie Lee Charitable Foundation, Fong Shu Fook Tong Foundation, and Croucher Foundation.

## AUTHOR CONTRIBUTIONS

JFH, ZZ designed the experiments; ZZ, DL, XL, WS conducted the electrophysiological and behavioral experiments in Figs. 2-5; CXZ, YP, ML, PT conducted the electrophysiological and behavioral experiments in Figs. 6-7; YPG, XC collected and analyzed the data in Fig.1; HTW, LLH collected and analyzed the data from cultured neurons; HMF, CXZ, YJP, XL, ZDW, KW, ML collected the data of behavioral experiments; RJ, CXZ, JTB, XS, XC collected and analyzed the anatomy data; and FQX prepared some experimental setup; JFH, ZZ, and XC wrote the manuscript.

## AUTHOR INFORMATION

The authors declare no competing financial interests.

## METHODS

### Animals

Sprague Dawley rats with clean external ears were used for behavioral study with no surgery (Fig.1) and for pre-experimental sessions in which they were trained to use 2 tones as cues for reward retrieval (Figs. 2-4). Rats were surgically implanted with electrodes and cannulae, and those that successfully finished the behavioral experiments were included in the study. Both male and female (8-10-week old) C57/BL/6 (C57, wild-type), *CCK-ires-Cre* (Cck^tm1.1(Cre)Zjh/J^, C57 background), and CCK-CreER (Cck^tm2.1(Cre/ERT2)Zjh/J^, C57 background, CCK^-/-^) mice were used for immunohistochemistry, *in vivo* extracellular recordings, and behavioral experiments (Figs. 5-6). Animals were confirmed to have clean external ears and normal hearing and were housed in a 12-hour-light/12-hour-dark cycle. All procedures were approved by the Animal Subjects Ethics Sub-Committees of City University of Hong Kong and The Hong Kong Polytechnic University.

### Auditory and visual stimuli

Auditory stimulus (AS) were digitally generated using a computer-controlled TDT Auditory Workstation and delivered through a coupled electrostatic speaker (EC1, TDT). Sound pressure level was calibrated with a condenser microphone (Center Technology, Taipei, Taiwan). Pure tones of 60-70 dB SPL were used to screen rats in the pre-training experiments. The visual stimulus (VS) was white light generated by light-emitting diodes placed 5 cm above the center hole of the behavioral apparatus. When the light was on, the illumination at the bottom of the apparatus was 26 Lux.

### Association of auditory and visual stimuli

Experimental rats (n=7) were exposed to 10 pairings of a combined auditory/visual stimulus (AS-VS) with footshock. Control rats (n=8) were presented with 10 pairings of just the VS with footshock. Conditioning of animals in the experimental and control groups was confirmed by freezing in response to a presentation of the combined AS-VS or VS alone, respectively, (both without footshock).

In the purely behavioral experiment (Fig.1), the experimental group was exposed to 10 presentations of a compound conditioned stimulus (CS) consisting of an AS (1800-ms white noise) and VS (2000-ms light). The auditory and visual stimuli were arranged serially such that the onset of the VS preceded the onset of the AS by 200ms and the 2 stimuli co-terminated. The VS 200ms was placed prior to the AS because it took time for the VS to reach to the AC and it might be easier to attract the animal’s attention to both stimuli in a serial presentation. Each CS presentation was followed immediately by footshock (600ms; 0.5-0.9mA; fear conditioning apparatus was homemade). The inter-trial interval was variable (30 or 60s). The AS was omitted in the control group.

Two days after the classical conditioning, both groups were trained on the subject-initiated sound-reward protocol^11^. After water intake was regulated, rats were presented in a custom-made cage with an array of 3 parallel holes^38^. An infrared sensor in the center hole detected nose-poking. On each trial rats nose-poked in the center hole to initiate the trial, then were presented with the sound. Another infrared sensor in the rightmost hole detected whether the rat’s arrival was within 2s of the onset of the sound. If it was within 2s, water reward was delivered, otherwise, water was not delivered. (The leftmost hole was not used in the present study.)

To make sure that the rats used the AS as a cue for reward availability, rats were required to keep their nose in the central hole for a varying interval of 100-1200ms before the AS was triggered. Training occurred in 5 stages: (1) rats were trained to obtain water from the rightmost hole; (2) rats were required to nose-poke in the central hole to receive a water reward in the right hole; (3-5) rats were trained to keep their nose in the central hole for varying amounts of time before reward-reception. The holding time varied randomly from 200 to 800ms (Stage 3), 400 to 1500ms (Stage 4), and 100 to 1200ms (Stage 5). Rats moved from one stage to the next when they had successfully obtained the reward in 4 successive trials. In Stage 5, the performance criterion was successful reward retrieval in 9 successive trials. One rat did not complete Stage 3 within 1h and was excluded from the study.

Because both control and experimental rats initiated the trials and retrieved the reward after hearing the AS, we termed this the “subject-initiated sound-reward protocol.”

On the day following mastering the subject-initiated sound-reward protocol, rats were tested in the same general paradigm except that the AS was combined with a VS. We reasoned that association between the VS and the AS would lead the rat to continue to perform the task successfully. We termed this the “subject-initiated light-reward (SiLR) protocol.” We categorized the SiLR protocol trials as successful or unsuccessful if the rat triggered the light and moved to the rightmost hole for the reward or if the rat triggered the light but failed to move to the rightmost hole for the reward, respectively. We computed the success rate as the number of successful trials divided by the number of successful and unsuccessful trials.

To monitor the progress of different stages of training, the following parameters were monitored: the time interval between the poking of the center and reward holes, the reaction time that was defined as the time interval between the cue presentation and arrival of the head in the reward hole, and the time difference between the above 2.

A successful trial was defined as one in which a rat retrieved the reward after the VS was triggered. Only trials in which the rat nose-poked long enough to trigger the VS were included in the statistical analysis.

### Surgery

For the experiments in Figs. 2-5, rats were anesthetized with sodium pentobarbital (50mg/kg i.p.; Ceva Sante Animale Co., Libourne, France), and anesthesia was maintained at a dose of 15mg/kg/h. Atropine sulphate (0.05mg/kg, s.c.) was administered 15 min before anesthesia to inhibit tracheal secretions. A local anesthetic (xylocaine, 2%) was liberally applied to the incision site. Animals were prepared for surgery as previously described^11,39^. Rats were mounted in a stereotaxic device, and a midline incision was made in the scalp after liberal application of a local anesthetic (2% xylocaine). Bilateral craniotomies were performed over the temporal lobe (3.0-6.5 mm posterior, 3.0-5.0 mm ventral to bregma) to access the AC, and the dura matter was opened. Body temperature was maintained at 37-38°C with a heating blanket.

Before electrode implantation, tungsten microelectrodes with impedances of 1-3MΩ (Frederick Haer & Co., Bowdoinham, ME) were used to identify the AC. Electrodes were positioned with an oil hydraulic micromanipulator controlled from outside the soundproof room. Neuronal signals recorded by the microelectrode, together with auditory and visual signals, were amplified and stored using Tucker-Davis Technologies (OpenEX, TDT, Alachua, FL) and Axoscope software (Axon Instruments, Sunnyvale, CA).

We then implanted guide cannulae for drug infusion and homemade electrode arrays which typically consisted of 6 electrodes (one stimulating electrode, 4 recording electrodes, and one reference electrode) into the AC of each hemisphere. Stimulating electrodes were made of insulated stainless steel wire (A-M Systems, Carlsborg, WA) with an impedance of <100kΩ. Reference and recording electrodes were made of insulated tungsten wire (California Fine Wire Co., Grover Beach, CA) with an impedance of 0.5-1MΩ. The electrode array was held by a micromanipulator and penetrated into the cortex. Electrodes in the 2 hemispheres were deliberately implanted symmetrically at slightly different locations to avoid invoking strong commissural connections. Ground electrodes for stimulation and recording were separately connected to screws on the skull. The electrodes were advanced to a depth of 1000-1100µm where neurons showed stable auditory responses, and the skull opening was covered with a layer of silicone (World Precision Instruments, Sarasota, FL). The connection sockets of the electrodes were cemented to the skull with the cannulae. Rats were then housed in their home cages and recovered for 5 days prior to experiments.

For experiments in Fig.6–7, wild-type, CCK-ires-Cre and CCK^-/-^ mice were anesthetized with pentobarbital sodium (0.8mg/kg, i.p.) and anesthesia was maintained throughout surgery and neuronal recordings with periodic supplements. Atropine sulphate (0.05mg/kg, s.c.) was administered 15min before induction of anesthesia to inhibit tracheal secretions. Briefly, animals were mounted in a stereotaxic device, and a midline incision was made in the scalp. A craniotomy was performed at the temporal lobe (2-4mm posterior, 1.5-3mm ventral to bregma) to access the AC, and the dura mater was opened. In the experiment in which virus injection was applied to the entorhinal cortex, another craniotomy was performed (4-5mm posterior, and 2.5-3.5mm lateral to bregma).

### Auditory cortex stimulation and reward retrieval

Five days after surgery, rats underwent behavioral training, during which they learned to retrieve a water reward cued by perceiving EAC. Water was restricted to 50% of normal intake prior to training. Rats were placed in a homemade cage with 3 horizontally aligned holes with infrared sensors and were first manually guided to poke their noses into the center hole before moving to the left or right hole, where a drop of water (15-20µl) was delivered. Rats then underwent formal training consisting of 4 stages. In Stage 1, a nose-poke in the center hole triggered a high- or low-frequency sound stimulus, which indicated specific hole for reward retrieval (i.e., for 7 rats, high-or low-frequency indicated the left or right hole, respectively; vice versa for the other rats (see Video 5). In Stage 2, rats were required to perceive EACL or EACR (5 pulses at 80-150µA, 20Hz) delivered after a nose-poke in the center hole. Pulses were generated by the TDT system and delivered through the stimulating electrode (of low impedance) via isolator (ISO-Flex, A.M.P., Israel). Stimulation of the left AC indicated reward availability in the right hole, and stimulation of the right AC indicated reward availability in the left hole. Infrared sensors detected whether rats arrived at the correct hole within 5s from the onset of stimulation, in which case a drop of water was delivered. Stimulations were delivered to only one hemisphere in one session (100 trials), and switched to another hemisphere in another session until a 90% success rate was reached. In Stage 3, stimulations were delivered to one hemisphere to another hemisphere for 10 trials each. Rats were required to reach a success rate of 90% or higher. In Stage 4, stimulations were delivered to either hemisphere in a pseudo-random manner. Training finished when rats reached a success rate of 90% or higher (see Video 6).

### Pairing the VS with EAC in anesthetized rats in the presence of CCK

After training, rats underwent a baseline testing phase during which a VS (white light, 500-ms duration and 26-Lux illumination) was randomly presented at a very low probability (10 out of 1000 trials). Behavioral responses to the light were recorded. No water reward was delivered with the presentation of the VS. Responses to 10 presentations of the VS were collected (except for one rat that received only 3 presentations).

Rats were then anesthetized with sodium pentobarbital. AC neuronal responses to the VS were measured before cholecystokinin octapeptide (CCK-8, abbreviated as CCK, 0.5-1.0µl, 10ng/µl, 0.1µl/min, Tocris) was infused into the left or right AC. Within 15 min after infusion, VS was then paired with EAC in the targeted hemisphere (2 pulses at 80-150µA, 20Hz). In each stimulation, VS (500ms) was presented, followed by EAC at the offset of VS. Forty pairings (10-s inter-stimulation-interval in between) occurred on each day for 5-6 days. Neuronal responses to the VS in both the targeted and naïve hemispheres were recorded.

After fully recovery from anesthesia, further post-intervention tests of behavioral responses to the VS were conducted. Responses were assigned as “Decision Index.” For example, when the left hole was the hole that was “engineered” to be associated with reward, approaching the left hole was scored 1; showing a tendency to approach the left hole but stopping before reaching the hole was scored 0.75; showing no obvious tendency to approach either hole was scored 0.5; showing a tendency to approach the right hole but stopping before reaching the hole was scored 0.25; and approaching the right hole was scored 0. When the right hole was the “engineered” hole, scores were assigned vice versa. To check whether a visuoauditory association had been established in awake rats, animals were placed in the apparatus while all holes were temporally blocked, and VS was presented for 30 times and the corresponding neuronal activities were recorded.

### Vehicle controls

To eliminate the possibility of non-specific effects of infusion or other manipulations of rats during anesthesia, we designed 2 control experiments. First, we infused CCK into one hemisphere and artificial cerebrospinal fluid (ACSF) into the other hemisphere at the same time, and paired the VS with simultaneous EAC of both hemispheres. Second, we infused ACSF into one hemisphere and performed stimulus pairings. One hour later, we infused CCK into other hemisphere and repeated the stimulus pairings.

### Optogenetic stimulation of auditory neurons and pairing with VS

AAV-CaMKIIa-ChR2-mCherry (10^13^gc/ml, 1µl, UNC vector core) was infused into rat’s primary cortex (5.0mm posterior, 4.0mm ventral to bregma). 4 weeks later, animal was anesthetized with pentobarbital sodium, and the response of auditory neurons to 472nm laser, AS and VS was examined to confirm the readiness of these neurons in the pairing procedure. CCK was then infused into auditory cortex, and 500-ms VS was presented for 30 trials, each of which was followed by a 5-ms laser pulse at the end. The inter-trial-interval was 10s. After pairing, auditory neurons’ response to VS was monitored for 10min.

### Artificial association between auditory and electrical stimuli induced by HF laser stimulation

In the experiment shown in Fig.6, AAV-DIO-ChR2-mCherry (10^13^gc/ml, UNC vector core) was infused into the entorhinal cortex of CCK-ires-Cre or CCK^-/-^ mice (4.3mm posterior, 3.2mm lateral, and 4.5mm ventral to bregma). Tamoxifen (150mg/kg, i.p.) was injected for 5 consecutive days, started from 4 days after virus injection. Mice recovered for at least 6 weeks to allow the virus to infect entorhinal cortical neurons. An optical fiber with 4 electrodes (~200kΩ, 300µm between electrodes, California Fine Wire Co., Grover Beach, CA, USA) was implanted into the AC. Neuronal responses to laser stimulation of AC, the AS, and the VS in field potential and unit responses were recorded by the implanted recording electrodes.

Single pulse EAC was used as a conditioned stimulus in the cued fear conditioning protocol. Initially, 5 pulses (0.5ms, 5Hz, 100-150µA) were applied. The onset delay between the first pulse and the footshock (0.5s, 0.5mA) was 5s. The number of pulses was gradually reduced to one based on mouse performance. After mice showed more than 2s of freezing within the first 5s of a single pulse, the VS as used before was used to test baseline of freezing. We then anesthetized mice and recorded baseline fEPSPs in response to the VS. High-frequency (HF; 80Hz, 5 pulses, 5-ms duration, 10mW) or LF (1Hz, 5 pulses) laser stimulation was delivered to the AC, followed 1s later by 5 pairings of the VS and single pulse EAC at 1Hz. The onset of the VS was 100ms earlier than the onset of EAC. This stimulation protocol was repeated 4 times with a 10-s interval. fEPSPs in response to the VS were recorded for 15 min before and after the pairing. A second pairing was performed 15min after the first pairing, and fEPSPs were recorded for another 15min. Neuronal response to the VS that were 3 standard deviations above or below means were excluded. Mean fEPSP slopes before and after the pairings were calculated by liner regression and analyzed using two-way ANOVA. Freezing responses to the VS and EAC were measured 24-48h after the pairing and analyzed using one-way ANOVA, separately.

### Immunohistochemistry

CCK-ires-Cre and CCK^-/-^ mice injected with AAV-DIO-ChR2-mCherry were fully anesthetized by an overdose of pentobarbital sodium and perfused with 30ml cold phosphate-buffered saline (PBS) and 30ml 4% paraformaldehyde. Brain tissue was removed, post-fixed, and treated with 30% sucrose at 4ºC for 2 days. Brain tissue was sectioned on a cryostat (40µm) and preserved with antifreeze buffer (20% glycerin and 30% ethylene glycol diluted in PBS) at -20ºC. Sections were incubated with anti-Dsred (Takara, 632496, Rabbit 1:500) multiclonal antibody at 4ºC for 48 h. After 4 times PBS washing, sections were incubated with secondary antibody (Jackson, 488, 111302) for 1.5h at 37ºC. All the sections were mounted on slides and subsequently imaged (20× magnification) by using an LSM 710 confocal microscope (Zeiss).

### Associative Learning Test after CCK Receptor Antagonist Application in the AC

After the same anesthesia and surgery as mentioned earlier, a drug infusion cannula was implanted in each hemisphere of the AC of the C57 mouse. The mouse was allowed for recovery for 2 days. For the experimental group, the VS and AS were first presented in pairs to the mouse for 25 trials in each session after the auditory cortex was bilaterally infused with CCKB receptor (CCKBR) antagonist (L365, 260, 10µM in 2% DMSO-ACSF, 0.5µl in both sides, injection speed 0.05µL/min), and for 4 sessions each day in 3 days. For the control group, CCKBR antagonist was replaced with ACSF. On day 4, a baseline test for the percentage of freezing over a time period of 10s was carried out after the VS and AS were presented separately. The AS was then conditioned with footshock for 3 trials. On day 5, post-conditioning tests were carried out to the VS and AS separately. The freezing percentages of different groups to VS were compared by two-way ANOVA.

### Associative Learning Test after tetropeptide CCK (CCK-4) administration in CCK^-/-^Mice

A drug infusion cannula was implanted on the top of the venous sinus (transverse sinus) of the CCK^-/-^ mouse to administer drugs though i.v. infusion. After the i.v. injection of CCK-4 (CCK-4 Group; 0.01ml; 0.17µM) or saline (Saline Group; 0.01ml), the VS and AS were immediately presented in pairs to the mouse for 25 trials in each session for 4 sessions each day for 3 consecutive days. The rest of the procedure was the same as the previous experiment. To induce a similar percentage of freezing of C57 mice, the CCK^-/-^ mice needed 9 trials.

### Quantification and statistical analysis

Neuronal and behavioral responses were recorded simultaneously using a computer. Single-unit spikes were distinguished using spike sorting software (OpenSorter, TDT). We considered 3 standard deviations (SDs) above baseline as the threshold to distinguish spikes. K-means clustering method in OpenSorter was adopted to sort single-unit spikes. A unit with the largest amplitude and normal overlaid spike profile was chosen from each electrode. Another criterion was that the number of spikes with an inter-spike interval of <2ms in the histogram should be <0.2% of the total number of spikes. The timing of spike occurrence relative to stimulus delivery was calculated using MATLAB software. Peristimulus time histograms (PSTHs) were calculated over a bin size of 20ms for AC responses and 50ms for visual responses.

Because there was no guarantee of recordings from the same unit across days, we used Z-scores (mean ± standard error) to characterize neuronal responses and compare responses across different time points^38^. Z-scores of neuronal responses to visual stimuli within a certain time period were calculated against the mean spontaneous firing rate within the same period, thereby representing the distance between the neuronal responses and the mean of spontaneous firing in units of SD (Z = (x-µ)/δ; where x is the neuronal response in each trial, and µ and δ are the mean and SD, respectively, of spontaneous firing rates across all trials). Higher Z-scores typically indicate a larger neuronal response, although they also differ depending on the total number of testing trials. Changes in Z-score after each conditioning session were used to assess the effectiveness of conditioning to induce neuronal plasticity. Paired student’s *t*-tests were used to compare neuronal responses with spontaneous neuronal activity. One-way repeated measures analysis of variance (ANOVA) was used to test for differences in mean Z-scores before and at different times after stimulus pairing sessions. Tukey’s post-hoc tests were used for mean comparisons. Statistical significance was set at *p*<0.05.

## SUPPLEMENTARY MOVIES

**Videos 1-4**. Behavioral response to VS and AS in the one-choice operant conditioning task.

**Videos 5-6**. Behavioral response to AS and EAC in the two-choice water retrieval task.

**Videos 7 and 9**. Behavioral response to VS before stimulus pairing.

**Videos 8 and 10**. Behavioral response to VS after stimulus pairing.

**Videos 11-14**. Behavioral response to VS and AS before and after HF stimulus pairing.

**Videos 15-18**. Behavioral response to VS and AS before and after LF stimulus pairing

**Videos 19-26**. Behavioral response to VS and AS before and after fear condition in the C57 mice injected with CCK antagonist or ACSF.

**Videos 27-34**. Behavioral response to VS and AS before and after fear condition in the CCK^-/-^ mice injected with CCK or Saline.

## SUPPLEMENTARY FIGURE LEGENDS

**Supplementary Figure 1.**
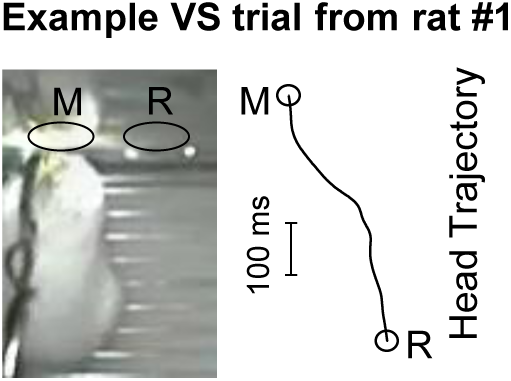
An exemplary head movement to the VS.

**Supplementary Figure 2.**
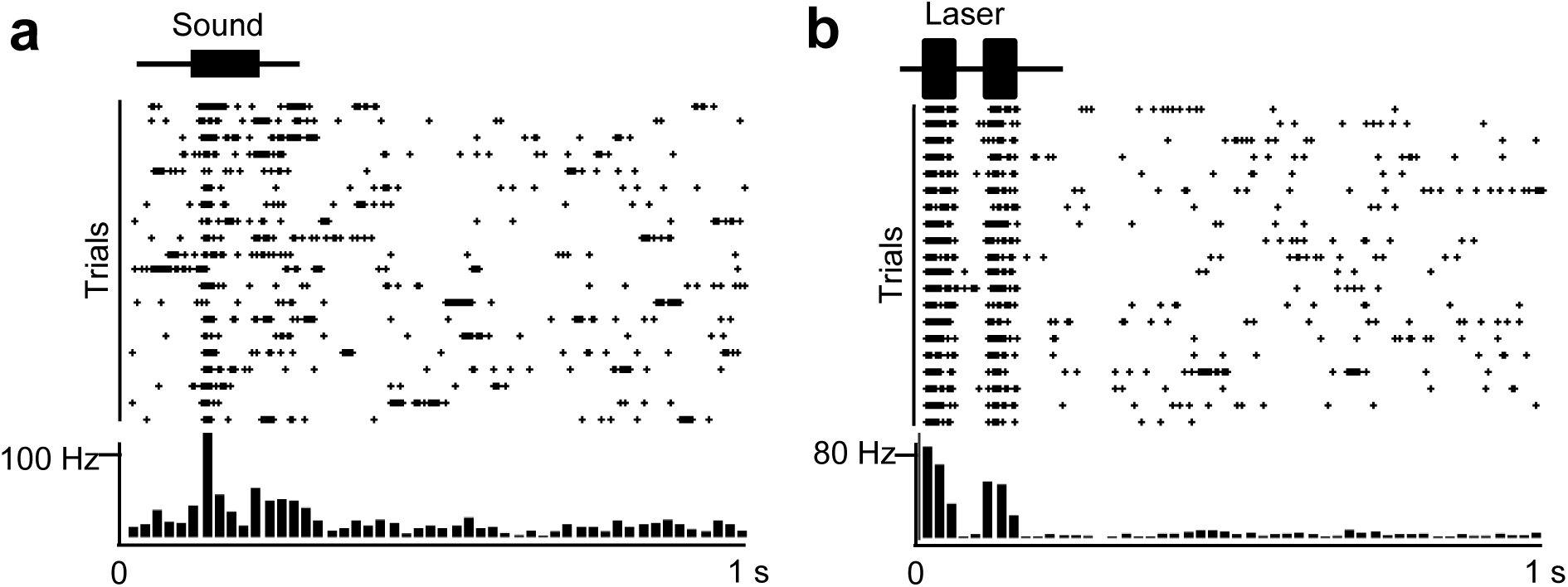
Neuronal response to AS (**a**) and laser (**b**) in the AAV-CaMkIIa-ChR2-mCherry injection AC.

## REFERENCE

1. L. R. Squire, S. Zola-Morgan, The medial temporal lobe memory system. Science 253, 1380–1386 (1991).

2. C. B. Canto, F. G. Wouterlood, M. P. Witter, What does the anatomical organization of the entorhinal cortex tell us? Neural Plast 2008, 381243 (2008).

3. L. W. Swanson, C. Kohler, Anatomical evidence for direct projections from the entorhinal area to the entire cortical mantle in the rat. J Neurosci 6, 3010–3023 (1986).

4. W. B. Scoville, B. Milner, Loss of recent memory after bilateral hippocampal lesions. J Neurol Neurosurg Psychiatry 20, 11–21 (1957).

5. S. Corkin, Lasting consequences of bilateral medial temporal lobectomy: Clinical course and experimental findings in HM. Sem. Neurology 4, 249–259 (1984).

6. E. Teng, L. R. Squire, Memory for places learned long ago is intact after hippocampal damage. Nature 400, 675–677 (1999).

7. S. H. Wang, C. M. Teixeira, A. L. Wheeler, P. W. Frankland, The precision of remote context memories does not require the hippocampus. Nat Neurosci 12, 253–255 (2009).

8. E. Lesburgueres et al., Early tagging of cortical networks is required for the formation of enduring associative memory. Science 331, 924–928 (2011).

9. K. S. Graham, J. R. Hodges, Differentiating the roles of the hippocampal complex and the neocortex in long-term memory storage: evidence from the study of semantic dementia and Alzheimer’s disease. Neuropsychology 11, 77–89 (1997).

10. L. R. Squire, R. E. Clark, B. J. Knowlton, Retrograde amnesia. Hippocampus 11, 50–55 (2001).

11. X. Chen et al., Encoding and retrieval of artificial visuoauditory memory traces in the auditory cortex requires the entorhinal cortex. J Neurosci 33, 9963–9974 (2013).

12. R. B. Innis, F. M. Correa, G. R. Uhl, B. Schneider, S. H. Snyder, Cholecystokinin octapeptide-like immunoreactivity: histochemical localization in rat brain. Proc Natl Acad Sci U S A 76, 521–525 (1979).

13. R. S. Greenwood, S. E. Godar, T. A. Reaves, Jr., J. N. Hayward, Cholecystokinin in hippocampal pathways. J Comp Neurol 203, 335–350 (1981).

14. C. Kohler, V. Chan-Palay, The distribution of cholecystokinin-like immunoreactive neurons and nerve terminals in the retrohippocampal region in the rat and guinea pig. J Comp Neurol 210, 136–146 (1982).

15. X. Li et al., Cholecystokinin from the entorhinal cortex enables neural plasticity in the auditory cortex. Cell Res 24, 307–330 (2014).

16. B. D. Winters, T. J. Bussey, Transient inactivation of perirhinal cortex disrupts encoding, retrieval, and consolidation of object recognition memory. J Neurosci 25, 52–61 (2005).

17. N. Suthana et al., Memory enhancement and deep-brain stimulation of the entorhinal area. N Engl J Med 366, 502–510 (2012).

18. J. F. Rehfeld, Immunochemical studies on cholecystokinin. II. Distribution and molecular heterogeneity in the central nervous system and small intestine of man and hog. J Biol Chem 253, 4022–4030 (1978).

19. P. Y. Deng et al., Cholecystokinin facilitates glutamate release by increasing the number of readily releasable vesicles and releasing probability. J Neurosci 30, 5136–5148 (2010).

20. T. Tsutsumi et al., Suppression of conditioned fear by administration of CCKB receptor antagonist PD135158. Neuropeptides 33, 483–486 (1999).

21. Y. Horinouchi et al., Reduced anxious behavior in mice lacking the CCK2 receptor gene. Eur Neuropsychopharmacol 14, 157–161 (2004).

22. M. Fendt, M. Koch, M. Kungel, H. U. Schnitzler, Cholecystokinin enhances the acoustic startle response in rats. Neuroreport 6, 2081–2084 (1995).

23. P. W. Frankland, S. A. Josselyn, J. Bradwejn, F. J. Vaccarino, J. S. Yeomans, Activation of amygdala cholecystokininB receptors potentiates the acoustic startle response in the rat. J Neurosci 17, 1838–1847 (1997).

24. S. A. Josselyn et al., The CCKB antagonist, L-365,260, attenuates fear-potentiated startle. Peptides 16, 1313–1315 (1995).

25. C. M. Lo et al., Characterization of mice lacking the gene for cholecystokinin. Am J Physiol Regul Integr Comp Physiol 294, R803–810 (2008).

26. S. Ramirez et al., Creating a false memory in the hippocampus. Science 341, 387–391 (2013).

27. T. J. Teyler, P. DiScenna, The hippocampal memory indexing theory. Behav Neurosci 100, 147–154 (1986).

28. J. O’Keefe, J. Dostrovsky, The hippocampus as a spatial map. Preliminary evidence from unit activity in the freely-moving rat. Brain Res 34, 171–175 (1971).

29. K. M. Igarashi, L. Lu, L. L. Colgin, M. B. Moser, E. I. Moser, Coordination of entorhinal-hippocampal ensemble activity during associative learning. Nature 510, 143–147 (2014).

30. D. Osipova et al., Theta and gamma oscillations predict encoding and retrieval of declarative memory. J Neurosci 26, 7523–7531 (2006).

31. D. B. Headley, N. M. Weinberger, Gamma-band activation predicts both associative memory and cortical plasticity. J Neurosci 31, 12748–12758 (2011).

32. G. A. Calvert et al., Activation of auditory cortex during silent lipreading. Science 276, 593–596 (1997).

33. R. A. Andersen, Multimodal integration for the representation of space in the posterior parietal cortex. Philos Trans R Soc Lond B Biol Sci 352, 1421–1428 (1997).

34. S. Higuchi, Y. Miyashita, Formation of mnemonic neuronal responses to visual paired associates in inferotemporal cortex is impaired by perirhinal and entorhinal lesions. Proc Natl Acad Sci U S A 93, 739–743 (1996).

35. S. Nabavi et al., Engineering a memory with LTD and LTP. Nature 511, 348–352 (2014).

36. M. D. Whim, P. E. Lloyd, Frequency-dependent release of peptide cotransmitters from identified cholinergic motor neurons in Aplysia. Proc Natl Acad Sci U S A 86, 9034–9038 (1989).

37. D. Shakiryanova, A. Tully, R. S. Hewes, D. L. Deitcher, E. S. Levitan, Activity-dependent liberation of synaptic neuropeptide vesicles. Nat Neurosci 8, 173–178 (2005).

## References

38. G.H. Otazu, L.H. Tai, Y. Yang, A.M. Zador, Engaging in an auditory task suppresses responses in AC. Nat Neurosci 12, 646–654 (2009).

39. X.J. Yu, X.X. Xu, S.G. He, J. He, Change detection by thalamic reticular neurons. Nature Neurosci 12, 1165–1170 (2009).

